# Coordinated regulation of osmotic imbalance by c-di-AMP shapes ß-lactam tolerance in Group B *Streptococcus*

**DOI:** 10.1101/2024.03.27.586946

**Authors:** Terry Brissac, Cécile Guyonnet, Aymane Sadouni, Ariadna Hernandez Montoya, Elise Jacquemet, Rachel Legendre, Odile Sismeiro, Patrick Trieu-Cuot, Philippe Lanotte, Asmaa Tazi, Arnaud Firon

## Abstract

*Streptococcus agalactiae* is among the few pathogens that have not developed resistance to ß-lactam antibiotics despite decades of clinical use. The molecular basis of this long-lasting susceptibility has not been investigated, and it is not known whether specific mechanisms constrain the emergence of resistance. In this study, we report the conserved role of the signaling nucleotide cyclic-di-AMP in susceptibility to ß-lactams, demonstrating that inactivation of the phosphodiesterase GdpP in *S. agalactiae* confers ß-lactam tolerance. Characterization of the c-di-AMP signaling pathway reveals antagonistic regulation by the transcriptional factor BusR, which is activated by c-di-AMP and negatively regulates ß-lactam susceptibility through the BusAB transporter and AmaP/Asp23 cell envelope stress complex. Furthermore, we show that the simultaneous inhibition of osmolyte transporters activity and transcription by c-di-AMP has an additive effect, sustaining ß-lactam tolerance. Finally, we expanded the analysis of ß-lactam tolerance using random transposon mutagenesis, uncovering a convergent pattern of mutations involving the KhpAB small RNA chaperone and the S protein immunomodulator. Overall, our results demonstrate that c-di-AMP acts as a turgor pressure rheostat, coordinating an integrated response to cell wall weakening due to ß-lactam activity, and identify mechanisms that may foster antibiotic resistance in *S. agalactiae*.

## INTRODUCTION

*Streptococcus agalactiae* (Group B *Streptococcus*: GBS) is the leading cause of bacterial invasive infection during the first three months of life ^1,2^. Infection of the newborn mainly occurs vertically during parturition in case of colonization of the mother vaginal tract. Prenatal screening and intrapartum antibiotic prophylaxis (IAP) decrease bacterial burden and risk of invasive infection during the first week of life ^3^. Beta-lactam antibiotics, especially penicillin G and amoxicillin, remain the first-line antibiotics for IAP and early neonatal infections. Despite decades of selective pressure, no ß-lactam resistant GBS isolates have been identified to date. Similarly, the closely related opportunistic pathogen *Streptococcus pyogenes* (Group A *Streptococcus*: GAS) remains sensitive to ß-lactams despite their widespread use to treat common respiratory tract infections. The scientific and clinical communities have long been puzzled by this fortunate situation ^4^. Are we only lucky or does specific biological processes hamper the emergence and selection of ß-lactam resistance in ß-hemolytic streptococci?

The recent isolation of non-susceptible ß-lactam GBS clinical isolates, mostly documented in Japan and US, suggests that we may end up with no standardized therapeutic options ^5,6^. The non-susceptible GBS isolates have increased minimal inhibitory concentrations (MICs), which remains below or close to the clinical susceptibility breakpoint. The reduced susceptibility is due to mutations in the *pbp2x* gene encoding the main target to which penicillin G binds ^7^, as similarly observed in non-susceptible GAS isolates ^8^. Mutations in a penicillin-binding protein (PBP) are often a first step towards resistance, which requires additional mutations to compensate for their fitness cost. For instance, isolation of non-susceptible ß-lactam *Streptococcus pneumoniae* isolates was rapidly followed by the emergence of resistant strains, a process facilitated by the pneumococcal competence machinery ^9–11^.

In addition to PBP mutations and acquisition of non-susceptible PBP variants or ß-lactamases, additional mechanisms contribute to ß-lactam resistance in pathogenic species. Especially, mutations in the GdpP phosphodiesterase have been recently described in *S. pneumoniae* and *mec*-negative *Staphylococcus aureus* clinical isolates with low-level ß-lactam resistance ^12–16^. The GdpP enzyme hydrolyzes cyclic-di-AMP (cdA), a signaling nucleotide acting as second messenger. The main cellular function of cdA is to regulate turgor pressure ^17^. Remarkably, the function of cdA in osmotic homeostasis is conserved in Firmicutes, but the precise molecular mechanisms have evolved, even in closely related species ^18,19^. This is probably the consequence of an ancient and essential regulatory mechanism that has had time to adapt to the specific environment and physiology of each species.

The synthesis of cdA is closely linked to the synthesis of the cell wall, which is essential for resisting turgor pressure and maintaining cell integrity ^20,21^. The essential enzyme GlmM, which synthesizes an early cell wall metabolite, interacts with and inhibits the cdA cyclase DacA, while the corresponding genes are co-transcribed from a conserved operon ^22–24^. However, cdA itself does not appear to directly regulate cell wall synthesis. The cdA mode-of-action is essentially based on the regulation of transporters which control intracellular concentrations of potassium and zwitterionic osmolytes (*e.g.*, trimethylglycine also called betaine). In addition to directly modulating transporter activities, cdA also regulates their expression by more species-specific mechanisms involving either two-component systems ^25,26^, transcriptional regulators ^27–29^, or riboswitches ^30,31^. Osmotic homeostasis is primarily achieved by osmolytes transporters but is also coupled to metabolic and physiological responses ^32^. In some species, cdA signaling evolves to additionally regulate the tricarboxylic acid cycle ^33,34^ and the general stress response mediated by the Rel enzyme synthesizing the alarmone (p)ppGppp ^35–38^.

Alterations in cdA metabolism have been associated with changes in ß-lactam susceptibility in several species, including laboratory and clinical isolates, but the underlying mechanism remains unclear ^29,39–43^. In this study, we first investigated whether cdA has a conserved role in susceptibility to ß-lactams in the almost universally susceptible GBS pathogen. Then, we untangled the cdA signaling pathway and show that osmolyte transporters individually contribute to ß-lactam susceptibility and that their co-regulation by cdA, directly and at the transcriptional level via the BusR regulator, confers ß-lactam tolerance. A genome-wide screen of ß-lactam decreased susceptibility confirms the role of cdA signaling and identifies additional pathways, including a conserved RNA binding protein with a predicted Hfq-like sRNA chaperone activity. This study reveals the molecular basis of cdA-dependent ß-lactam tolerance and prompts an in-depth surveillance and characterization of mechanisms that can promote the development of resistance in pathogens that are still susceptible to this precious class of antibiotic.

## RESULTS

### Inactivation of the c-di-AMP phosphodiesterase GdpP confers ß-lactam tolerance

We first investigated whether the second messenger cdA had a conserved role in ß-lactam susceptibility in a species which has always remained clinically susceptible. In GBS, cdA is synthesized from two ATP molecules by the cyclase DacA and hydrolyzed into pApA by the phosphodiesterase GdpP ^27^. DacA, and thus cdA synthesis, is essential for growth unless the concentration of osmolytes is tightly limited in the growth medium or compensatory mutations allow growth in usual media. ^27^. To test the susceptibility to ß-lactams, we therefore used a Δ*gdpP* mutant with a high cdA intracellular concentration made in the NEM316 WT strain ^27^. Firstly, conventional antibiotics susceptibility testing shows similar MICs for the Δ*gdpP* mutant and the WT control, whether by broth microdilution or by gradient tests using Etests (Fig. 1A).

**Figure 1.**
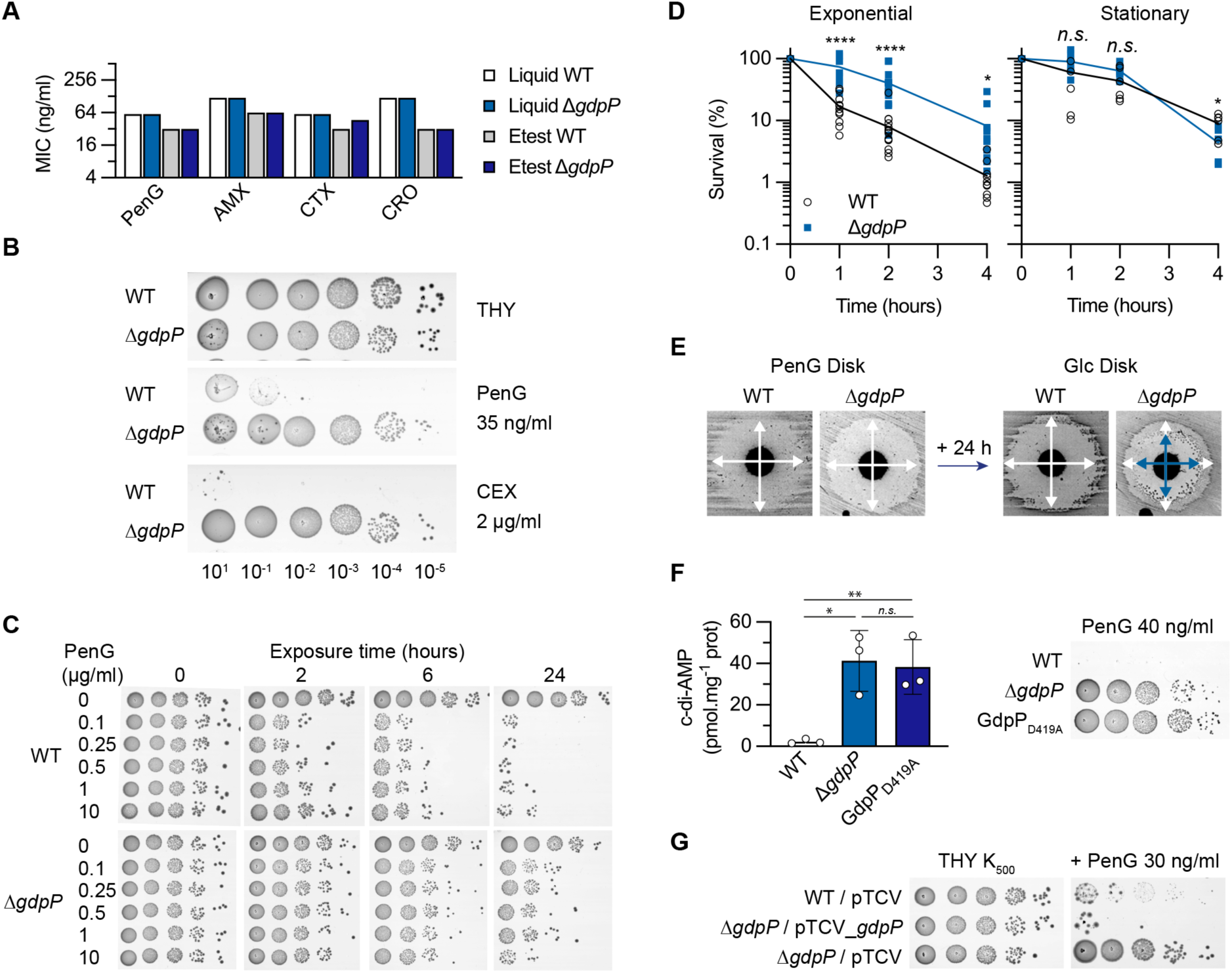
Inactivation of the cyclic-di-AMP phosphodiesterase GdpP confers beta-lactam tolerance. **(A)** Minimum inhibitory concentration (MIC) of antibiotics. Conventional tests in liquid (broth microdilution) and solid (gradient Etest strips) in MH-F media with the WT strain (white and grey bars) and the Δ*gdpP* c-di-AMP phosphodiesterase deletion mutant (light and dark blue bars) for penicillin G (PenG), amoxicillin (AMX), and the third-generation cephalosporin cefotaxime (CTX) and ceftriaxone (CRO). **(B)** Spotting assays of the WT strain and Δ*gdpP* mutant. Overnight cultures were serially diluted (10^1^ to 10^-5^) and spotted on the surface of THY agar supplemented with PenG or the first-generation cephalosporin cephalexin (CEX). Images were taken after 16-24 hours incubation at 37°C in aerobic condition with 5% CO_2_. **(C)** Dose and time-dependent killing of the WT strain and Δ*gdpP* mutant. High PenG concentrations (0.1 – 10 µg/ml) are added to exponential growing cultures of the WT strain and Δ*gdpP* mutant. Aliquots were taken at the indicate time, serial diluted, and spotted on THY without antibiotic. **(D)** Time-killing of exponential and stationary phase cultures of the WT strain and Δ*gdpP* mutant in presence of 10 µg/ml PenG. Aliquots were taken at the indicate time and dilution were plated on THY for colony-forming unit numeration. Data are from 7 and 6 independent experiments for exponential and stationary time, respectively. Unpaired t-test were used to compare the WT and the mutant at each time point (* p < 0.05; **** p < 0.0001). **(E)** Adaptation of the TD-test for GBS antibiotic tolerance. Diluted cultures of the WT strain and Δ*gdpP* mutant are spread on MH media and PenG disks (10 µg) were added. After a 24h incubation, PenG disks were removed and replaced by Glucose (Glc) disks (10 mg). Crossed white arrows highlight inhibition zones after 24 hours, and crossed blue arrows highlight the decrease in the inhibition zone of the Δ*gdpP* mutant after an additional 24h incubation with the Glc disk. **(F)** Penicillin tolerance depends on the GdpP phosphodiesterase activity. Left panel: quantification of intracellular c-di-AMP by LC-MS in the WT strain, the Δ*gdpP* mutant, and the catalytic inactivated GdpP_D419A_ mutant. Data represent means and SD calculated from biological triplicate (N = 3) and analyzed with unpaired t-test (* p < 0.05; ** p < 0.01). Right panel: spotting assays on PenG plate. **(G)** Genetic complementation of the Δ*gdpP* mutant. Spotting assay with the WT and the Δ*gdpP* mutant containing the empty (pTCV) or complementing (pTCV_*gdpP*) vectors. Kanamycin (K_500_) is added to maintain the selective pressure for the vectors.

In contrast, spotting assays reveal growth of the Δ*gdpP* mutant on THY plates containing concentrations of penicillin G and cephalexin that inhibit the WT strain (Fig. 1B). Time-killing experiments with high penicillin concentrations (MIC x 2 < 0.1 to 10 µg/ml < MIC x 200) added to exponentially growing cultures before spotting on THY plates without antibiotic show that the Δ*gdpP* mutant is killed more slowly than the WT strain (Fig. 1C). Noteworthy, the bactericidal activity is initially inversely proportional to the drug concentration when exceeding four times the MIC, a phenomenon known as the “Eagle effect” and first described in the 1940s for GBS ^44^, for the WT control only (Fig. 1C). Time-killing experiments using CFU counts confirmed the advantage of the Δ*gdpP* mutant, which exhibited a decreased killing rate compared to the WT strain (Fig. 1D). Slow killing kinetics and no difference between the Δ*gdpP* mutant and the WT strain were observed when stationary phase cultures were used (Fig. 1D), in agreement with the mode of action of ß-lactams requiring active cell wall metabolism and division. Adaptation of the TD-test, a modified disk-diffusion assay for antibiotic tolerance ^45,46^, further shows that the Δ*gdpP* mutant can survive lethal concentration of penicillin (Fig. 1E).

To determine whether tolerance is a result of GdpP enzymatic activity, we substituted the conserved catalytic residue D_419_ ^20,47^ with an alanine by targeted mutagenesis in the WT strain. The GdpP_D419A_ catalytic-minus and the Δ*gdpP* mutants have similar high intracellular cdA concentration and ß-lactam phenotype (Fig. 1F). In addition, complementation of the Δ*gdpP* mutant with a vector containing a WT copy of *gdpP* (pTCV_*gdpP*) abolishes the phenotype of the Δ*gdpP* mutant on penicillin-containing plates (Fig. 1G). Overall, a high intracellular concentration of cdA confers ß-lactam tolerance, characterized by similar MICs but slow killing kinetics ^48^, demonstrating that cdA is involved in the response of GBS to ß-lactams.

### cdA catabolism impacts physiology, cell envelope and division

Antibiotic tolerance is frequently dependent on the growth rate and/or metabolic activity of the target cell, especially for ß-lactams ^49–51^. At first glance, the individual growth curves in rich medium (THY) show a fitness defect in the exponential phase for the Δ*gdpP* mutant compared with the WT strain, with a doubling time of 27.2 +/- 1.1 and 34.4 +/- 0.5 minutes, respectively (Fig. 2A). However, preliminary microscopic observations revealed significant heterogeneity and major morphological defects in the Δ*gdpP* mutant. Closer examination by scanning and transmission electron microscopy show morphological defects affecting the ultrastructures of the cell surface, both membranous and parietal (Fig. 2B). Instead of chains of dividing cocci, Δ*gdpP* cells are heterogenous, often swollen, with rough surface and septation defects. Cell envelopes are irregularly thickened, cells are frequently compartmentalized by multiple curved septum-like structures, and invaginated membranous structures are also observed (Fig. 2B and 2C). The WT morphology is restored in the Δ*gdpP* mutant with the complementing *gdpP* vector but not with the empty vector (Fig. 2D). Overall, the absence of *gdpP* leads to severe morphological alterations, with large effects on cell envelope and division.

**Figure 2.**
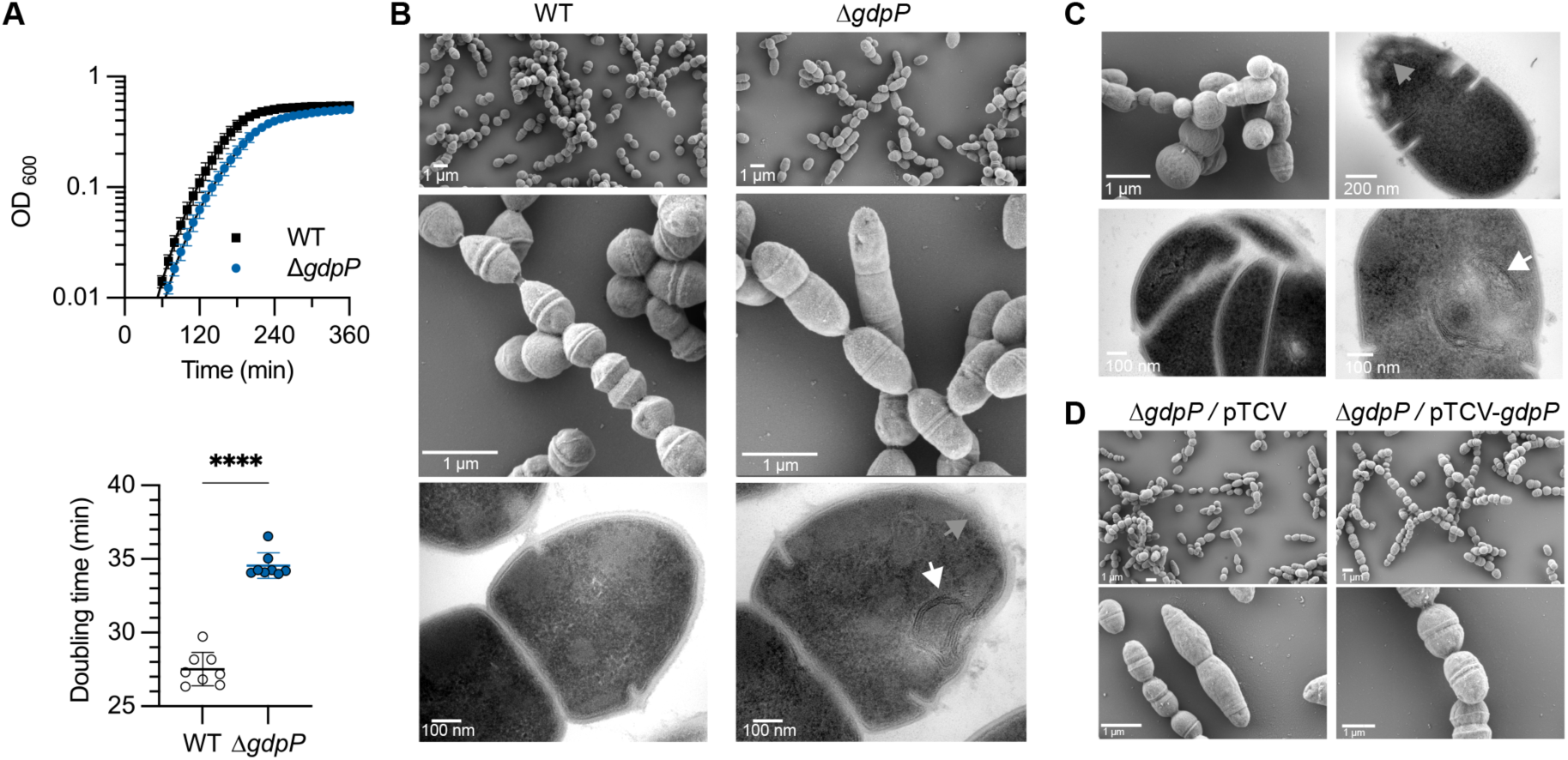
Cyclic-di-AMP phosphodiesterase deficiency leads to morphological and cell envelope defects. **(A)** Growth curves (upper panel) and doubling time in exponential phase (bottom panel) of the WT strain and Δ*gdpP* mutant in THY at 37°C. Means and SD are calculated from biological replicates (N = 8) and analyzed using unpaired t-test (**** p < 0.0001). **(B)** Scanning and transmission electronic microscopy of the WT strain and Δ*gdpP* mutant at similar scales. The white and grey arrows highlight intracytoplasmic membrane structures and areas with cell envelopes of heterogeneous thickness, respectively. **(C)** Additional electronic microscopy illustrating the heterogeneity and atypical ultrastructure of Δ*gdpP* cells. **(D)** Representative images of the Δ*gdpP* mutant with the empty (pTCV) or complementing (pTCV_*gdpP*) vectors observed by SEM.

### The BusR transcriptional repressor antagonizes ß-lactam tolerance

The morphology of the Δ*gdpP* cells suggests large-scale perturbations involving, either directly or indirectly, cell wall synthesis. To characterize the associated stress responses, we analyzed the transcriptome of the Δ*gdpP* mutant in exponential growth phase at 37°C in rich media. In total, 69 genes are significantly differentially expressed (DEG : |log_2_ FC| > 1 and p-adj < 10^-4^), among which 28 are part of multicopy integrative genomic elements known as TnGBS ^52^ that we have excluded for the rest of the analysis (Fig. 3A and Supplementary Table S1A). Of the remaining DEG, 11 and 30 are up- and down-regulated, respectively. Except for two uncharacterized genes, fold changes and significance are relatively minor (Fig. 3A), and analysis of gene function does not provide a clear pattern (Supplementary Table S1A). To gain confidence, we analyzed the transcriptome of a second Δ*gdpP* mutant made in a different WT background (strain BM110, of the same capsular serotype III and belonging to the hypervirulent clonal complex CC-17 ^53^). Applying the same thresholds, 67 DEG were identified including 39 in prophages (Fig. 3A and Supplementary Table S1B). Comparative analysis between the two backgrounds reveals only 22 genes with a conserved significant differential expression (p-adj < 10^-4^), among which only 10 with a conserved threshold above |log_2_ FC| > 1 (Fig. 3B and Supplementary Table S1C and S1D). Overall, transcriptomes suggest a strain-specific adaptation to the absence of *gdpP*, but do not allow to unambiguously identify a conserved response.

**Figure 3.**
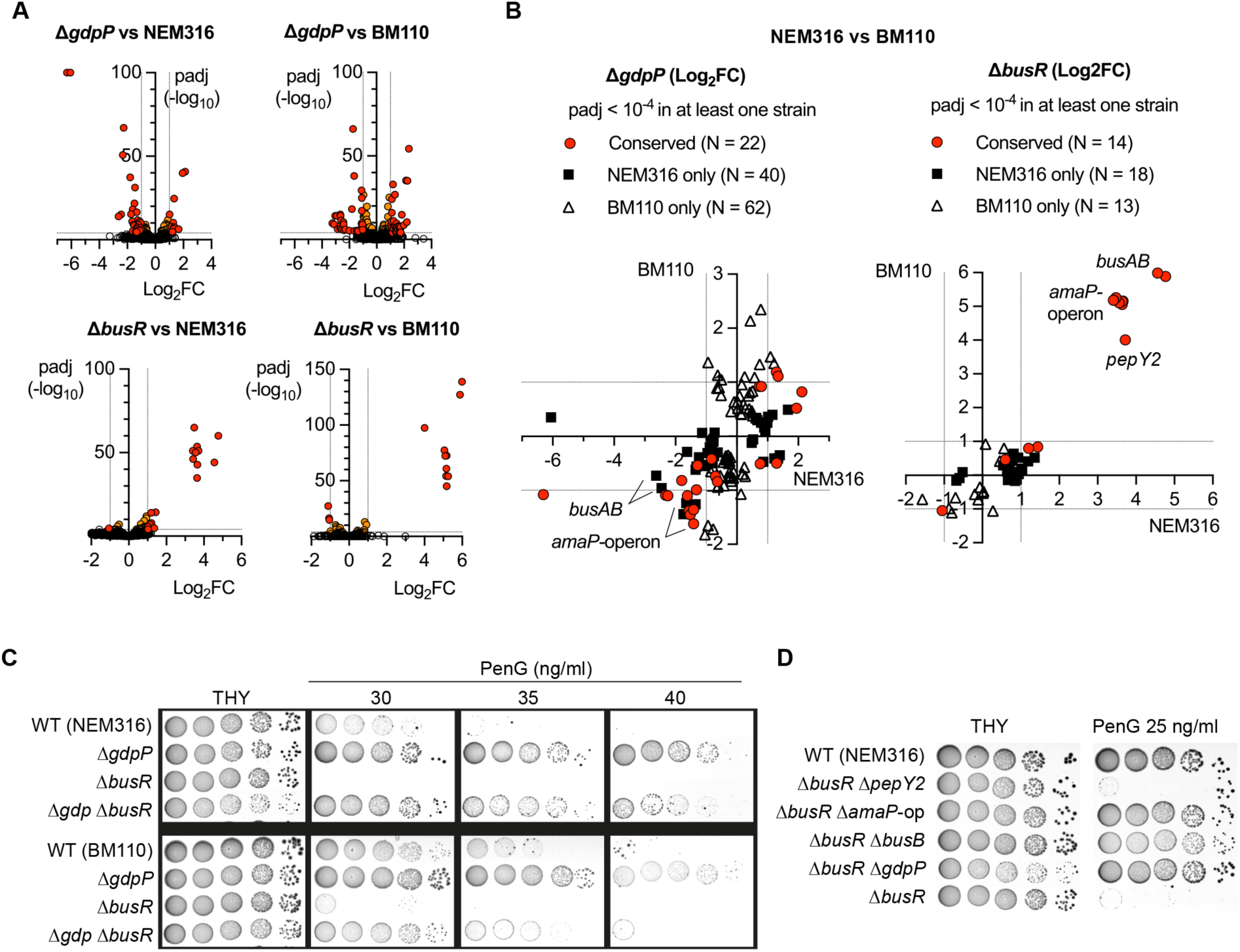
ß-lactam tolerance depends on the c-di-AMP-activated BusR repressor. **(A)** Transcriptome analysis by RNA-seq of Δ*gdpP* and Δ*busR* mutants in two WT backgrounds (NEM316 and BM110). Colored dots on volcano plots highlight significant differential expression (p-adj < 10^-4^) with fold changes |FC| > 2 (red dots) and |FC| < 2 (orange dots). **(B)** Comparative analysis of the Δ*gdpP* and Δ*busR* transcriptomes between the two WT backgrounds. Red dots highlight conserved statistical significance (p-adj < 10^-4^) in the two strains, and black squares and white triangles highlight statistical significance in NEM316 only or BM110 only, respectively. **(C)** Tolerance of Δ*gdpP* to ß-lactam depends on the cdA-activated BusR repressor. Spotting assays (10^-1^ to 10^-5^ serial dilution) of Δ*gdpP*, Δ*busR*, and the double Δ*gdpP* Δ*busR* mutants in the two WT backgrounds on THY supplemented with penicillin. **(D)** Susceptibility of Δ*busR* to ß-lactam depends on *busB* and *amaP* operon. Spotting assays with Δ*busR*, and the double Δ*busR* Δ*pepY2*, Δ*busR* Δ*amaP-*operon, Δ*busR* Δ*busB*, and Δ*busR* Δ*gdpP* mutants in the NEM316 background.

Despite the relatively low level of information provided by RNA-seq analysis, we and others have previously shown that the BusR transcriptional regulator is allosterically activated by cdA and directly represses the *busAB* operon encoding a betaine transporter ^27–29,54^. Consistently, *busAB* genes are among the most repressed in the Δ*gdpP* mutants but reach statistical significance in the NEM316 background only (Fig. 3B). This indicates that the BusR repressor is active in the WT strains and over-activated by the high cdA concentration in the Δ*gdpP* mutants. We therefore deleted *busR* in the WT strains and in their corresponding Δ*gdpP* mutants to test ß-lactam susceptibilities. Deletion of *busR* is associated with a fitness cost, especially in the Δ*gdpP* backgrounds (Supplementary Fig. S1), indicating that BusR must be tightly regulated and that its inactivation is antagonistic with a high cdA concentration. Interestingly, Δ*busR* mutants are more susceptible to penicillin compared to their WT parental strains (Fig. 3C). Moreover, deletion of *busR* in the Δ*gdpP* mutants decreases the growth advantage of the parental Δ*gdpP* mutants on penicillin-containing plates (Fig. 3C). The Δ*gdpP* phenotype is thus dependent on the transcriptional repressor BusR, which is already active in the WT strain and negatively regulates genes conferring ß-lactam susceptibility.

### The BusR signaling pathway regulates ß-lactam susceptibility

To characterize the BusR regulon beyond the known direct repression of the *busAB* operon, we analyzed the transcriptome of the Δ*busR* mutants in the two WT backgrounds (Fig. 3A and 3B). In addition to *busAB* (FC = 23 to 64-fold, 10^-45^ < p-adj < 10^-140^), BusR negatively regulates a 7-genes operon thereafter called the *amaP-*operon (FC = 10 to 37-fold, 10^-35^ < p-adj < 10^-78^) adjacent to, and translated in the same direction as, the *busR* operon (Supplementary Fig S1 and Supplementary Table S1E). The seven small proteins (65 to 194 amino acids) encoded in the *amaP*-operon are predicted to form a membrane-localized protein complex with redundant functional subunits (3 x Asp23-domain proteins PF04226/PF03780/DUF322 including AmaP, 2 almost identical GlsB-like proteins PF04226, 1 x DUF2773, 1 x CsbD-like family PF05532). In addition, one monocistronic gene (*pepY2*, FC = 13 to 16-fold, 10^-52^ < p-adj < 10^-98^) encoding a predicted transmembrane protein containing two PepSY ectodomains (PF03413) is also highly significantly repressed by BusR (Fig. 3B and Supplementary Table S1E).

The three genetic units (*busAB*, *amaP*-operon, *pepY2*) define the BusR regulon, which is distinct from the indirect transcriptional perturbations observed in individual Δ*busR* transcriptomes (Fig. 3B). Putative BusR binding sites are detected close to *amaP*-operon and *pepY2* transcriptional start sites suggesting direct BusR-repression (Supplementary Fig S1), as previously demonstrated for BusR on the *busAB* promoter ^54^. To test the role of the regulon in the susceptibility of the Δ*busR* mutant to ß-lactams, we deleted *busB*, the whole *amaP*-operon, and *pepY2* in the Δ*busR* mutant (*n.b.* in the NEM316 WT strain only). Deletion of *busB* and of the *amaP*-operon restores the growth of the parental Δ*busR* mutant on penicillin-containing plates (Fig. 3D). Overall, BusR negatively regulates three genetic units, including two that confer susceptibility to ß-lactams susceptibility when BusR is inactive: the betaine transporter BusAB and an AmaP/Asp23/GlsB complex with a probable function in cell envelope homeostasis ^55–57^. In the context of a Δ*gdpP* mutant with a high level of cdA, BusR is overactivated and the repression of the regulon contributes to the tolerance of the Δ*gdpP* mutant to ß-lactams.

### Joint regulation of osmolyte transporter activity and expression confers tolerance to ß-lactams

The opposite but additive ß-lactam phenotypes of Δ*gdpP* and Δ*busR* mutants reveal a mechanism based on cdA activation of the BusR transcriptional repressor. However, the cdA signaling network in GBS is a set of negative regulations (Fig. 4A) which also includes the direct inhibition of the potassium transporters KtrAB and TrkAH and of the zwitterionic transporter subunit OpuCA ^27^. In addition, cdA likely inhibits the RCK_C domain containing protein EriC, a chloride channel protein necessary to reestablish the ionic balance after osmolyte uptake ^27^. Lastly, binding of cdA to the small CBS domain protein CbpB abolishes the CbpB-Rel allosteric interaction leading to decreased (p)ppGpp synthase activity ^38^, which is a conserved mechanism of antibiotic tolerance ^58,59^.

**Figure 4.**
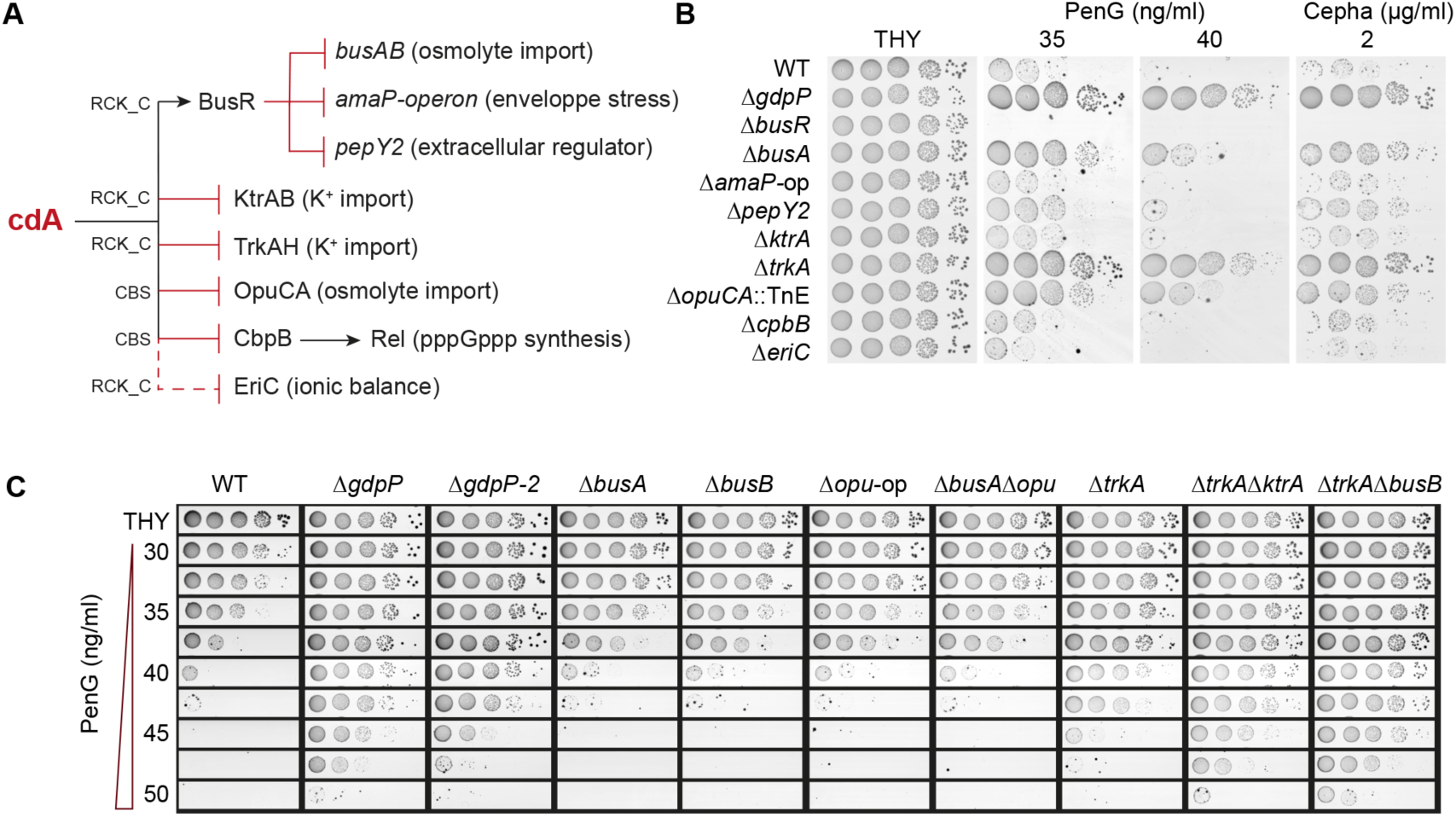
Coordinated regulation of osmolyte transporters confers ß-lactam tolerance. **(A)** Diagram of the c-di-AMP (cdA) signaling network in GBS. The cdA binding domain (RCK_C or CBS) of each effector is indicated on the connecting lines, with arrows indicating activation and final bars indicating repression or inhibition. The dotted line denotes predicted cdA binding to the RCK_C domain of EriC that has not been demonstrated experimentally. **(B)** Individual effectors contribute to ß-lactam susceptibility. Spotting assays (10^-1^ to 10^-5^ serial dilution) of a collection of deletion (Δ) or insertion (::TnE) mutants for each individual cdA effector on ß-lactam-containing plates. **(C)** Additive effect of cdA effectors on penicillin susceptibility. Spotting assays of single and double deletion mutants on plates containing increasing concentrations of penicillin, in increments of 2.5 ng/ml. An independent Δ*gdpP*-2 mutant was included for validation.

All the components of the signaling pathway are inhibited by the high cdA concentration in the Δ*gdpP* mutant. To test their individual contribution, we used a collection of deletion and insertional mutants in the WT strain (n.b. NEM316). On penicillin-containing plates, the Δ*busA*, Δ*opuCA::TnE*, and Δ*trkA* mutants grow on ß-lactam concentration higher than those inhibiting the WT strain, without reaching the phenotype of the Δ*gdpP* mutant (Fig. 4B). To test whether transporters have cumulative effects, we generated additional single deletion mutants (Δ*busA*, Δ*opu*-operon) and double deletion mutants for the two zwitterionic transporters (Δ*busA* Δ*opu*-op), two potassium transporters (Δ*ktrA* Δ*trkA*), and a pair of potassium-betaine transporters (Δ*trkA* Δ*busB*).

After sequencing the genome of all mutants (Supplementary Table S2A), we identified two issues. First, we unexpectedly identified two different NEM316 WT profiles, differing by 14 and 8 SNPs (5 in common) compared to the reference sequence (Supplementary Table S2B). The 14 SNPs profile was already reported ^60^ and the 8 SNPs profile corresponds to the oldest mutants generated in the laboratory (Δ*gdpP* ^27^ and Δ*opuCA::TnE* ^61^). Although the Δ*gdpP* phenotypes are complemented (Fig. 1 and Fig. 2) and out of cautious, we constructed a new mutant (Δ*gdpP*-2) in the 14 SNPs profile and validated the absence of any secondary mutation (Supplementary Table S2A). Second, the *pepY2* mutants have secondary mutations in *glsB* or *lysM* genes, which we cannot exclude as compensatory mutations (Supplementary Table S2C). Consequently, the non-involvement of *pepY2* in ß-lactam susceptibility is not conclusive at this stage.

Focusing on potassium and zwitterionic transporters, parallel phenotypic analysis reveals three degrees of penicillin susceptibility. First, inactivation of the BusAB or OpuCA zwitterionic transporters confers a slight advantage in the presence of penicillin (Fig. 4C). The phenotype is similar between single deletion mutants and is not additive (Fig. 4C). Second, inactivation of the potassium importer TrkA leads to a stronger phenotype closer to Δ*gdpP* mutants. Third, inactivation of either the BusAB or KtrA transporter in the Δ*trkA* mutant further increases the growth advantage of the Δ*trkA* single mutant to a level even slightly superior to the Δ*gdpP* mutant (Fig. 4C). These results show that the Δ*gdpP* ß-lactam tolerance is potentiated by the simultaneous inactivation of potassium and zwitterion importers. Notably, the stronger phenotype is associated with the inhibition of potassium uptake, which is the first cellular response in case of hyperosmotic stress. The contribution of the second cellular response, the uptake of zwitterion to compensate the deleterious effects of potassium, is dependent on the BusR regulator and is dampened by transporters redundancy. Altogether, this reflects the dynamics of osmolyte exchanges coordinated by cdA through direct inhibition (KtrA, TrkA, OpuCA) and transcriptional repression (*busAB* through BusR activation).

### cdA primes cells against penicillin-induced osmotic shock

We next sought to test the effect of potassium and zwitterions on the Δ*gdpP* ß-lactam phenotype. Interestingly, we observed a ‘mirror’ phenotype on synthetic minimal media containing only trace amount of potassium (Fig. 5A). As expected, the WT strain grows on minimal media and is inhibited by the addition of penicillin. At the opposite, the Δ*gdpP* mutant is unable to grow unless penicillin is added (Fig. 5A). The addition of betaine (5 mM) circumvents the need for penicillin and restores Δ*gdpP* growth, while the addition of potassium (0.1, 0.5, or 5 mM) has a marginal effect, and the two osmolytes do not influence the growth of the mutant in the presence of penicillin (Fig. 5A). Altogether, this suggests that the cdA-inhibition of high affinity osmolyte transporters inhibits the growth of the Δ*gdpP* mutant and that penicillin weakens the cell wall allowing osmolyte uptake.

**Figure 5.**
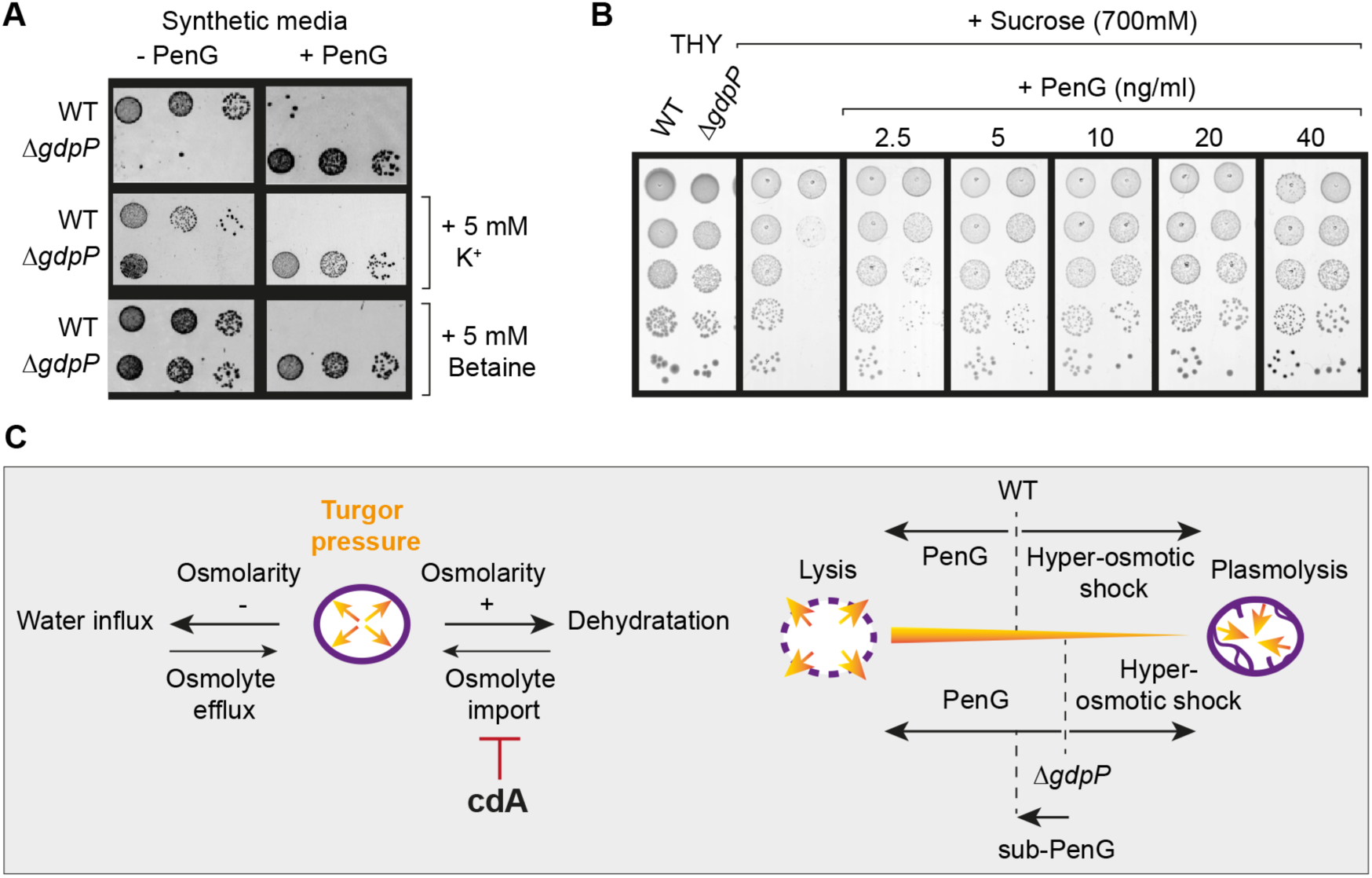
C-di-AMP balances osmotic and ß-lactam susceptibilities. **(A)** Penicillin alleviates Δ*gdpP* osmolyte requirements on minimal media. Spotting assays of the WT strain and Δ*gdpP* mutant on synthetic media supplemented with penicillin, potassium, and betaine at the indicated concentration. **(B)** Sub-inhibitory ß-lactam concentrations counteract Δ*gdpP* osmo-susceptibility. Spotting assays of the WT strain and Δ*gdpP* mutant under hyperosmotic condition (THY supplemented with sucrose) with and without penicillin. **(C)** Osmotic regulation by cdA and adaptation to osmotic imbalances. Left panel: the cellular turgor pressure (orange arrows) is mainly regulated through osmolytes exchanges to respond to change in osmolarity. The import of osmolytes is necessary to counteract cellular dehydration due to increased osmolarity and is inhibited by cdA. Right panel: the high cdA concentration in the Δ*gdpP* mutant results in a hyperosmotic stress physiology in a standard environment. The advantage against the bactericidal activity of ß-lactam is at the cost of sensitivity to hyper osmotic condition. Fluidification of the cell wall with ß-lactam sub-inhibitory concentration allows osmolyte exchange and re-establishes a WT-like turgor pressure.

The Δ*gdpP* mutants have been previously demonstrated to be susceptible to hyperosmotic stress in GBS and other species due to the inhibition of osmolyte transporters required to reestablishes the turgor pressure ^23,27,28,62^. To test whether cell wall rigidity contributes to the Δ*gdpP* osmo-susceptibility, we added sub-inhibitory concentrations of penicillin to high osmolarity growth medium. Strikingly, cell wall fluidization by sub-inhibitory concentrations restores the growth of the Δ*gdpP* mutant in hyperosmotic condition (Fig. 5B). Overall, this suggests that the Δ*gdpP* mutant is in a physiological state corresponding to hyperosmotic stress under normal growth conditions (Fig. 5C). The Δ*gdpP* mutant is unable to cope with standard hyperosmotic stress, but its reduced turgor pressure confers an advantage when challenged with the bactericidal activity of penicillin (Fig. 5C).

### Convergent mutations impact ß-lactam susceptibility

We currently have little information on ß-lactam tolerance, and obviously on resistance, in ß-hemolytic streptococci to compare cdA metabolism and regulation of osmotic homeostasis with other mechanisms ^63^. We therefore set-up a genome-wide screen for ß-lactam tolerance and reduced susceptibility in GBS. First, we established the condition of the screen by mixing WT and Δ*gdpP* cultures at different ratio and by plating dilution on penicillin-containing plates. Only the fraction of cell corresponding to the Δ*gdpP* cell number grows above 35 ng/ml of penicillin (Supplementary Fig. S2). Secondly, we engineered a modified transposon delivery system to construct a pool of 1.2×10^5^ insertional mutants in the NEM316 WT strain. Sequencing of the transposon-chromosome junction in 240 isolated colonies shows a random distribution of the transposon along the chromosome (Fig. 6A and Supplementary Table S3A).

**Figure 6.**
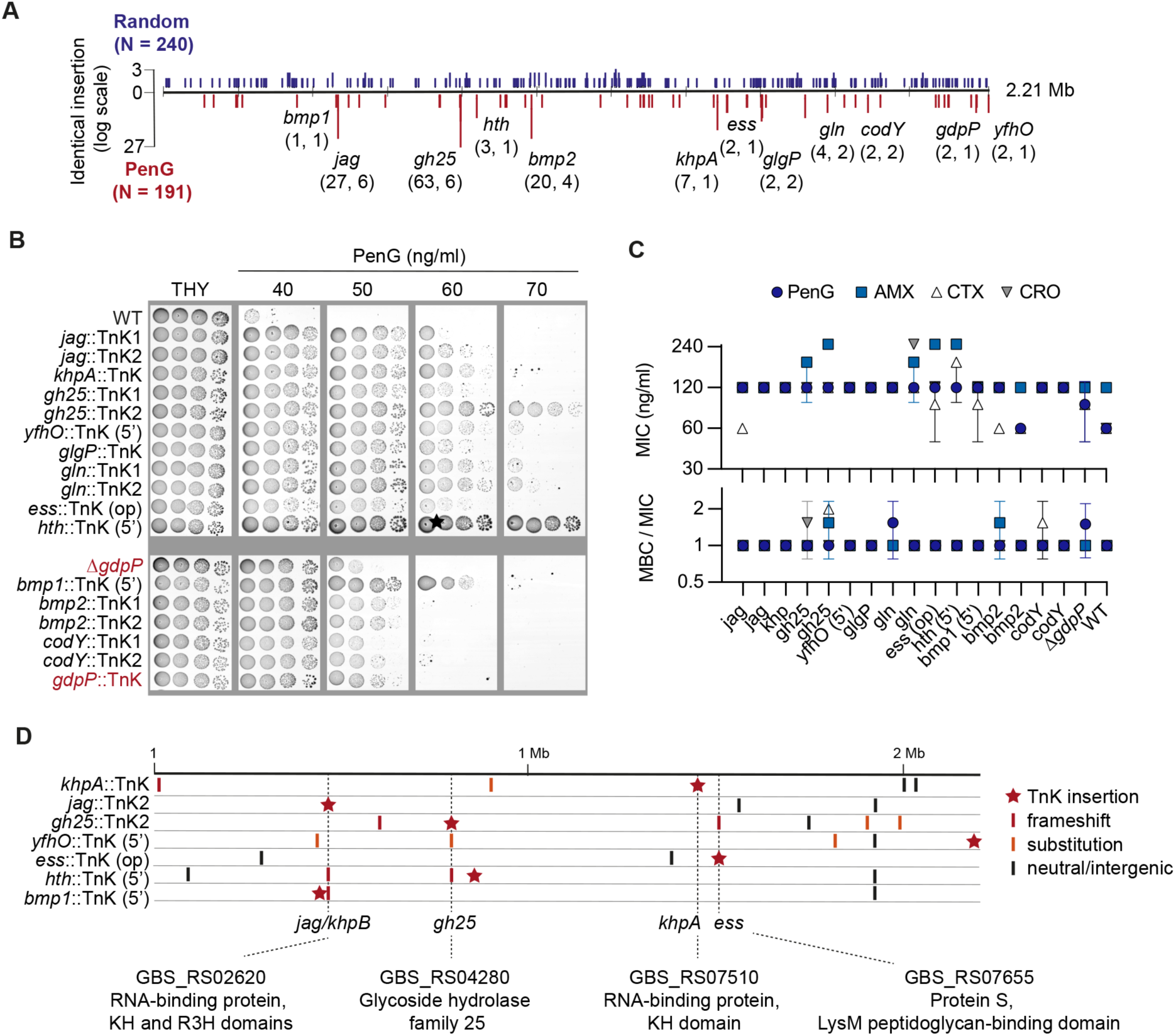
Genome-wide screening for ß-lactam tolerance in GBS. **(A)** Distribution of transposon insertions on the GBS chromosome. Vertical bars represent identical transposon insertions along the 2.1 Mb genome in randomly isolated colonies on THY (N = 240, upper blue bars) or THY supplemented with 35-40 ng/ml PenG (N = 191, lower red bars). Gene names are specified for mutants selected for validation. The numbers in brackets below the gene names indicated the total number of transposon insertions followed by the number of independent integration sites. **(B)** Validation of penicillin phenotypes. Spotting assay with selected insertional mutants (::TnK) on plates with increasing PenG concentration. Insertions localized in promoters are indicated after the mutant name (5’) as well as insertions localized in non monocistronic transcripts (-op). The Δ*gdpP* mutant and a *gdpP*::TnK insertional mutant were included for comparison (highlighted in red). **(C)** Minimal inhibitory concentration (MIC) and minimal bactericidal concentration (MBC) for ß-lactams. MIC are determined in liquid (MH-F media) by serial dilution for penicillin G (PenG) amoxicillin (AMX), cefotaxime (CTX) and ceftriaxone (CRO). MBC are determined by numeration of viable bacteria after 16 hours of contact with the antibiotic in MH-F. Data are mean and SD of two independent experiments (N = 2). **(D)** Convergent pattern of secondary mutations in insertional mutants. The transposon insertion site (red star) and SNPs (vertical bars) along the chromosome are shown for seven insertional mutants. Dotted lines highlighted the three genes showing insertional and secondary mutations in independent mutans, as well as the KhpA subunit interacting with Jag/KhpB.

We next conducted a genome-wide screen on penicillin containing plates with the 1.2×10^5^ random insertional collection. A 100-fold reduction of bacterial viability is observed at 30 ng/ml, with a further 10-fold reduction in viability per 5 ng/ml drug increment (Supplementary Fig. S2). Sequencing of the transposon-chromosome junctions in 191 colonies isolated from the 35 and 40 ng/ml screening plates shows a highly biased insertion of the transposon at specific loci (Fig. 6A), with 69% (132/191) of the insertions into 5 genes at 29 independent chromosomal positions (Supplementary Fig. S2 and Supplementary Table S3). The most frequently inactivated gene (63/191 = 33% of the insertions, at 6 independent position) encodes for a secreted cell-wall lytic enzyme with a predicted lysozyme activity (GH25, glycosyl hydrolase family 25, PF01183). The six independent *gh25*::TnK mutants grow better than the WT in the presence of penicillin, but the orientation of the transposon in the *gh25* gene has an impact on the strength of the phenotype (Supplementary Fig. S2). This GH25 hydrolase is a candidate for being the main peptidoglycan hydrolase required for the bactericidal activity of ß-lactams in GBS ^64,65^.

To further validate the screen, we selected 16 mutants with an insertion into 11 genes, based on the number of independent insertions and on functional annotation. We also added one of the two mutants with an insertion of the transposon into *gdpP*. All insertion mutants grew at concentrations of penicillin that inhibited the WT strain (Fig. 6B) and, as expected, the *gdpP*::TnK and Δ*gdpP* mutants have a similar phenotype. Remarkably, twelve insertional mutants grow on penicillin-containing plates at concentration that inhibits the *gdpP* mutants (Fig. 6B). All mutants have similar ß-lactams MIC, within a 2-fold factor of the WT MIC (Fig. 6C). Moreover, the minimal bactericidal concentrations (MBC) after 16 hours of contact with ß-lactams are similar to the MIC for all mutants, giving an MBC/MIC ratio close to 1 (Fig. 6C), far from the threshold defining clinical antibiotic tolerance (MBC/MIC ≥ 32).

Genome sequencing of 7 selected insertional mutants reveals a convergent pattern of secondary mutations in 4 genes: *gh25*, *ess*, and *khpB (jag) - khpA* (Fig. 6D). For instance, secondary mutations in *gh25* are present in the *hth*::TnK and *yfhO*::TnK insertional mutants (Fig. 6D and Supplementary Table S3). Similarly, the *gh25*::TnK2 mutant has a secondary mutation in *ess*, encoding the protein S with a LysM peptidoglycan binding domain involved in immune evasion and probably cell wall metabolism ^66,67^, independently identified in the TnK screen (Fig. 6). Finally, the two sub-units of a predicted small RNA chaperone (KhpA and Jag/KhpB, PF13083) have been independently identified in the TnK screen, with loss-of-function *jag/khpB* secondary mutations also present in two additional TnK mutants including one mutated in *gh25* (Fig. 6D and Supplementary Table S3). Although the precise KhpAB function remains to be characterized, this global RNA chaperone has a conserved role in ß-lactam sensitivity and/or cell wall synthesis as recently described in several Firmicutes ^68–70^. In conclusion, our non-exhaustive screen reveals convergent pathways towards ß-lactams tolerance or reduced susceptibility in GBS with initial insertional mutations promoting the selection of additional mutations between functionally interacting genes. Detailed analysis of Gh25, *ess*/protein S and KhpAB is now required to elucidate the conserved and specific mechanisms involved in ß-lactam susceptibility in GBS.

## DISCUSSION

The cyclic-di-AMP signaling nucleotide is an essential regulator of bacterial physiology ^17^ and has emerged as a key determinant of low-level ß-lactam resistance in pathogenic species ^12–16^. This study shows that cdA is a conserved regulator of ß-lactam antibiotic response in a historically clinically sensitive species. Mechanistic analysis of the cdA signaling network reveals an integrated pathway dedicated to the maintenance of osmotic homeostasis in which individual effectors contributes to the overall response. Our model can be applied generically but needs to consider the unique characteristics of each species.

Our results provide evidence that regulation of cell turgor is the main mechanism underlying cdA-related cell wall phenotypes, as originally suggested ^71^. Inactivation of GdpP leads to resistance or tolerance to ß-lactam antibiotics and, at the opposite, absence of cdA synthesis is associated with increased susceptibility ^71^. Inactivation of potassium uptake systems counteracts the increased susceptibility of *Lactococcus lactis* mutant unable to synthesize cdA, in agreement with a model linking ß-lactam susceptibility with impaired osmotic regulation ^42^. However, the inactivation of cdA metabolizing enzymes also have pleiotropic phenotypes related to growth rate, morphology, and transcriptome. For instance, inactivation of GdpP (called Pde1) in *S. pneumoniae* is not associated with fitness defect and cell size are reduced in acapsulated strains only ^16^, whereas electronic microscopy shows aberrant cell envelope ultrastructure in *S. aureus* ^13,72^. Phenotypic variability needs to be linked to the evolution of the cdA signaling network. While the regulation of osmotic homeostasis is conserved, each species has a unique set of cdA-regulated genes and proteins, whose functions are often redundant.

The core of the system is the coordinated regulation of osmolytes transporters at the transcriptional and post-translational levels. In GBS, ß-lactam tolerance of the *gdpP* mutant is recapitulated by the simultaneous inactivation of cdA-regulated osmolytes transporters, either repressed by BusR (BusAB) or by allosteric inhibition (KtrA, TrkA, OpuCA). The activity of ß-lactams weakens the cell wall, causing an osmotic imbalance. Rebalancing involves exchanges of osmolytes in an ordered flow of potassium and zwitternionic molecules. Functional redundancy ensures that the system remains functional in a variable environment, considering factors such as extracellular osmolyte concentration and physico-chemical characteristics ^73^. Cda acts as a rheostat on these redundant transporters and the high concentration in the *gdpP* mutant locks osmolytes exchanges. The resulting reduction in turgor pressure has an impact on cell growth, probably through partial plasmolysis, as suggested by electronic microscopy. The advantage against the lytic activity of ß-lactams comes at a cost of a decreased ability to adapt to hyper-osmotic environment. It is likely that the individual contribution of the transporters varies from one species to another, depending on their physiological and environmental lifestyle, but that their simultaneous inhibition conservatively regulates ß-lactam tolerance or resistance.

A major difference between species is the cdA-dependent transcription mechanisms. In GBS, *L. lactis*, and *Clostridioides difficile*, the key transcriptional regulator is the BusR repressor ^27,29,54,74^. In addition to the conserved regulation of the *busAB* transporter genes, the BusR regulon includes the *amaP*-operon and the *pepY2* gene. Homologues of AmaP and related Asp23 proteins in *S. aureus* and *Bacillus subtilis* are involved in envelope homeostasis, either acting at the level of the cell wall or cell membranes ^55–57^. Additionally, the membrane spanning PepY2 protein containing two PepSY ectodomains, originally predicted to inhibit extracellular proteases or glycosylases ^75^, has homologues (TseB, CopB) recently characterized as part of the elongasome complex regulating PBP activity ^76,77^. It is thus likely that BusR integrates osmoregulation with envelope homeostasis. Turgor pressure exerts mechanical forces on cell envelopes, affecting cell growth, membrane tension and cell wall remodeling ^32^. Membrane polarization and flow of cell-wall metabolite intermediates across membranes are critical for adjusting cellular activity to the osmolarity of the environment ^78^. Further studies are necessary to define in which pathway AmaP/Asp23 and PepY2 are involved, without excluding a direct function in cell wall remodeling. Determining the dynamics of the response is also necessary. Especially, the *amaP* operon mediated ß-lactam susceptibility when BusR is inactivated but does not have a significant effect when WT cells are challenged with ß-lactams. This should be contextualized with the strongest effect observed for potassium transporters, suggesting that BusR transcriptional regulation is a secondary response that restores a new equilibrium in turgor pressure.

Inactivation of *gdpP* has been proposed to be a first step in the evolution towards high-level ß-lactam resistance ^16^. From an ethical point of view, the selection of ß-lactam-resistant GBS is prohibited by the scientific community, as is the introduction of a ß-lactamase gene, to prevent any risk of dissemination. In this study, we have first tested available mutants ^27^ and extended the analysis to a random collection of insertional mutants, similarly generated in other laboratories ^79,80^ and as recently done in *S. pyogenes* ^63^. Comprehensive screening by Tn-seq in *S. pyogenes* mostly reveals the function of the ClpX chaperone and CcpA metabolic regulator in reduced ß-lactam susceptibility ^63^. Our screen with individual mutants positively selected in similar conditions identified three main proteins or protein complex whose functions are cell-wall related but uncharacterized to date: the lysozyme-like GH25 hydrolase, the immunomodulatory Protein S with a peptidoglycan binding domain ^66,67^, and the small RNA chaperone KhpAB which regulates ß-lactam susceptibility in several species ^68–70^. Patterns of secondary mutations in the insertional transposon mutants suggest functional links between the three, either as additive or compensatory mechanisms.

Overall, we report tolerance or reduced susceptibility to ß-lactam in GBS independent of PBP mutations, which could pave the way for resistance. Characterization of the cdA signaling network shows that tolerance is due to its function in coordinating osmotic pressure, a model that could be applied to related pathogens. Furthermore, we have identified cell wall related mechanisms that may contribute to the distinctive features of GBS, particularly the absence of any reported clinical resistance to date.

## MATERIAL AND METHODS

### Bacterial strains and culture conditions

The NEM316 (CC-23) and BM110 (CC-17) strains are clinical isolates of capsular serotype III with sequenced genomes available under the NCBI references RefSeq NC_004368 ^81^ and NZ_LT714196 ^53^, respectively. Strains were grown in Todd-Hewitt supplemented with 1% yeast extract and buffered with 50 mM HEPES pH 7.3 (THY) incubated at 37°C in static condition. Growth curves were done in 96 well microplates (Clear, Flat bottom, Thermo Scientific) with 150 µl of diluted overnight culture (1/500) by well. Optical density (OD_600_) was automatically recorded every 10 minutes with 1 minute agitation by cycle at 37°C (TECAN Infinite). Doubling times are determined by fitting non-linear regression with a Malthusian growth model (GraphPad Prism 10) in exponential phase (R^2^ > 0.99). Chemically Defined Medium (CDM) is prepared as described ^27^ from a 2-fold stock solution without glutamine and potassium. Glutamine (2 mM) and indicated concentration of potassium and betaine are added extemporaneously, altogether with 2-fold melted BactoAgar solution.

*E. coli* strains (TOP10, Invitrogen or XL1-blue, Stratagene) used for vector construction were grown in LB with appropriate concentrations of antibiotics (ampicillin 100 μg/ml, kanamycin 25 μg/ml, or erythromycin 150 µg/ml). For selection and propagation of vector in GBS, THY is supplemented with erythromycin (10 µg/ml) or kanamycin (500 µg/ml).

### Antibiotic susceptibility tests

Minimal Inhibitory Concentration (MIC) in liquid is done following EUCAST guidelines in Mueller-Hinton Fastidious culture media (MH-F, Becton Dickinson) media using custom AST Sensititre 96 wells plates (ThermoScientific) and 18h of incubation at 37°C. Minimal Bactericidal Concentration (MBC) is done by numerating the number of viable bacteria by colony-forming units after MIC determination. MIC on agar plates is done with Etest strips (BioMerieux) on MH-F agar uniformly inoculated with a standardized bacterial solution (5.10^8^ CFU/ml) according to EUCAST guidelines.

Spotting assays are done on THY agar supplemented with the indicated concentration of antibiotic. Stocks solution of Penicillin G (1 mg/ml) are aliquots and stored at -20°C for single used and THY plates containing penicillin are prepared and used the same day. Overnight cultures are serial diluted (10-fold factor: 10^-1^ to 10^-5^) in PBS and 4 µl of each dilution is spotted at the surface of THY plates with and without antibiotic. Incubation is at 37°C in aerobic condition with 5% CO_2_. Time-killing by spotting assay. Cultures in early logarithmic growth phase (0.3 < OD_600_ < 0.4) in THY are adjusted to 10^8^ CFU/ml and 0.9 ml are distributed in 96 deep wells plates. Sterile water or 10-fold concentrated penicillin are added (0.1 ml), homogenized (MixMate, Eppendorf), and plates are incubated at 37°C. At the indicated time, cultures in deep wells plates are homogenized (MixMate) and an aliquot (50 µl) is taken, serial diluted in PBS (10^-1^ to 10^-5^) in a 96 well plate and spotted on THY without antibiotic. All steps are done with multichannel pipettes to uniformize time of antibiotic exposure between samples.

Time-killing by CFU quantification. Penicillin was added (time O) in 10 ml of adjusted cultures (10^8^ CFU/ml) in early logarithmic growth phase in THY. Incubation was resumed at 37°C and aliquots were taken at the indicated times. Serial dilutions in PBS (10-fold factor) were spread on THY without antibiotic to quantify the number of viable bacteria at each time points. Survival is the ratio of CFU at time X against CFU at time O.

TD-Test for antibiotic tolerance. The original TD-test ^46^ was adapted for GBS as described for *Staphylococcus aureus* ^45^. First, a disk diffusion assay was performed on MH medium uniformly inoculated with diluted GBS cultures and with penicillin disks (0.1 to 10 µg). After overnight incubation at 37°C in 5% CO_2_ atmosphere, penicillin disks are removed and carefully replaced by sterile disks containing glucose (10 µg). Tolerance is manifested by bacterial re-grown into the initial inhibition zones after an additional incubation at 37°C in 5% CO_2_ atmosphere for 24 hours. **Bacterial genetic and genome sequencing**

All the mutants used in this study, together with the summary of genome sequencing, are described in Supplementary Table S2. Oligonucleotides and construction of vectors are detailed in Supplementary Table S4. Deletion vectors were constructed by splicing-by-overlap PCR using high-fidelity polymerase (Thermo Scientific Phusion Plus) and Gibson assembly into the pG1 thermosensitive shuttle vector as described ^27^. Final PCR products contain the desired mutations flanked by 500 bp of sequences homologous to the target loci and end with 25 bp of sequences complementary to the pG1 vector. After Gibson assembly, vectors are introduced in *E. coli* XL1 blue (Stratagene) with erythromycin selection. Vector inserts are validated by Sanger sequencing (Eurofins Genomics).

Vectors are introduced in GBS by electroporation. Transformants are selected at 30°C (pG1 permissive replication temperature) with erythromycin. Integration of the vector by homologous recombination at the targeted loci is selected by streaking transformants on THY supplemented with erythromycin and incubated at 37°C (non-permissive temperature) and further isolation of single colonies in the same condition. Loss of the chromosomally integrated vector occurs through subcultures (n = 3 to 5) in THY at 30°C without antibiotic selective pressure. Single colonies were tested (n = 24-48) for the loss of the vector (erythromycin susceptibility) and by discriminatory PCR (MyTaq HS - Bioline) with specific oligonucleotides to select mutant over WT genotypes. Genomic DNA were purified following manufacturer instruction for Gram-positive bacteria (DNeasy Blood and Tissue – Qiagen) and sequenced (Illumina sequencing at Core facility or Eurofins Genomics). High quality reads in Fastq were mapped against the reference genome (162-fold coverage mean) and analysed with Geneious Prime (2019.2.3 - Biomatters Ltd).

### c-di-AMP quantification

Cyclic-di-AMP concentration was determined by LC-MS/MS following company’s instruction (Biolog LSI) and as previously described for GBS ^27^. Bacterial cultures in exponential growth phase were pelleted, washed with PBS, resuspended in 300 µl of nucleotide extraction buffer (acetonitrile/methanol/water; 2/2/1), incubated 15 min on ice, heated 10 min at 95°C, and incubated for an additional 15 min on ice. Cells were lysed by mechanical sharing with 0.1 µM beads (Precellys Evolution, Bertin) and clear lysates were recovered after centrifugation at 4°C. Lysis was repeated two times on cell debris with 200 µl of extraction buffer each time. Clear lysates were pooled and incubated 16 hours at -20°C for protein precipitation. After centrifugation (20 min, 4°C), supernatants were recovered and the whole extract was evaporated to dryness (Eppendorf Concentrator). Dry samples were sent to Biolog LSI for nucleotide quantification by LC-MS/MS. For sample normalization, total protein concentration was determined in the initial bacterial cultures.

### Electronic microscopy

Bacteria were grown in THY at 37°C until early logarithmic growth phase, harvested by centrifugation, washed twice in PBS, and fixed by incubation in a solution of 4% paraformaldehyde and 1% glutaraldehyde in 0.1M phosphate buffer (pH 7.2) for 24 hours. After two-step washing in PBS, bacteria were postfixed in 2% osmium tetroxide for 1 hour and dehydrated in graded ethanol solutions. For scanning electron microscopy (SEM), samples were finally dried with hexamethyldisilazane (HMDS), coated with platinum by sputtering, and observed with a Zeiss Ultra Plus SEM (Microscopy Department, University of Tours). For transmission electron microscopy (TEM), samples were embedded in Epon resin and allowed to polymerize for 48 hours at 60°C. Ultrathin sections of 90 nm were obtained, deposited on EM gold grids, and stained with 5% uranyl acetate and 5% lead citrate, before observation using a JEOL JEM-1011 microscope (Microscopy Department, University of Tours).

### RNA sequencing and analysis

RNA purification, sequencing and analysis were done as described ^60^. Total RNA was purified from three independent cultures done on different days. The culture conditions are THY inoculated (1/50) with an overnight culture and incubated at 37°C in static condition until exponential growth phase (OD_600_ = 0.5). Bacteria are harvested by centrifugation (5 minutes, 4°C) and washed with 1 ml cold PBS containing RNA stabilization reagents (RNAprotect, Qiagen) before flash freezing and storage at -80°C. Total RNA are extracted after cell wall mechanical lysis with 0.1 µm beads (Precellys Evolution, Bertin Technologies) in RNApro reagent (MP Biomedicals), and purified by chloroform extraction and ethanol precipitation. After resuspension in water (Invitrogen), residual DNA is removed (TURBO DNase, Ambion), RNA concentrations are quantified with fluorescent dye (Qubit RNA HS, Invitrogen) and RNA qualities are validated by electrophoresis (Agilent Bioanalyzer 2100).

Depletion of rRNA (FastSelect Bacterial, Qiagen), libraries construction (TruSeq Stranded mRNA, illumine) and sequencing (NextSeq 500, Illumina) were done following manufacturer instructions. Single-end strand-specific 75 bp reads were cleaned (cutadapt v2.10) and mapped on the GBS genomes (Bowtie v2.5.1). Gene counts and differential expression were analysed using DESeq2 (v1.30.1) in R (v4.0.5) ^82^. Normalization, dispersion, and statistical tests for differential expression were performed with independent filtering. Raw p-values were adjusted for each comparison (Benjamini and Hochberg multiple tests) and adjusted p-value lower than 0.005 were considered significant.

### Transposon mutagenesis

A minimal mariner transposon was constructed by PCR with oligonucleotides containing the inverted repeat, modified to contain MmeI restriction sites, used to amplify the kanamycin resistant marker of the pTCV vector (Supplementary Table S4). The purified and digested PCR product was cloned between the EcoRI-BamHI restriction sites of the thermosensitive pG1 vector. A second PCR was done to amplify the Himar9 hyperactive transposase encoding gene under the control of a *gyrA* constitutive promoter. The PCR product was cloned between BamHI-PstI restriction sites to give the pG_TnK vector.

The pG_TnK vector is introduced in GBS by electroporation with erythromycin selection at 30°C (permissive temperature of replication for the vector). Transformants are isolated and cultured in the same condition. After an overnight culture, a starting culture is inoculated (1/25) in THY without antibiotic and incubated at 37°C (non-permissive temperature of replication for the vector) for 2 to 4 hours. Dilutions (5 to 10-fold) were spread on THY with kanamycin (500 µg/ml) and incubated at 37°C. To control for vector loss and estimate transposition frequency, dilutions were also spread on THY at 30 °C and 37°C (total CFU), and THY with erythromycin at 30°C (total CFU containing the vector) and 37°C (chromosomal integration or mutation of the vector). From four biological replicate, transposition frequency is between 1.5×10^-4^ and 5.9×10^-5^ with 88.1 to 94.4% of kanamycin resistant - erythromycin susceptible colonies corresponding to chromosomally integrated TnK and loss of the vector backbone.

A total of 176 THY kanamycin plates inoculated from 4 independent starting cultures and with 5×10^2^ to 10^3^ colonies after incubation at 37°C were used to constitute the library collection. Bacteria were gently recovered with 4 ml of THY by plate, pooled, centrifuged (10 min, 4°C), washed with THY, resuspend in 4 x 25 ml glycerol 20%, and stored at minus 80°C by aliquots of 1.5 ml. Single tube were used for numeration on THY kanamycin at 37°c and single colonies were picked to confirmed erythromycin susceptibility (n = 96; > 95% ery^S^). Genomic DNA of isolated colony were purified from 7.5 ml of culture (DNeasy Blood and Tissue – Qiagen) with an additional step of cell lysis with microbeads (Precellys Evolution). Genomic DNA are used as template for Sanger sequencing with a transposon specific primer (BAC protocol, Eurofins). Sequence reads were mapped against the transposon end and flanking sequences are then mapped against the GBS genome to identify the transposition integration site. For screening, THY were inoculated (1/100) with -80°C library stocks and incubated 1 hours at 37°C for recovery before spreading dilutions on freshly prepared THY plates containing increasing concentration of penicillin. Plates are incubated at 37°C in a 5% CO_2_ incubator and colonies were further isolated on THY before genomic DNA purification and sequencing of transposon-chromosome junction (Eurofins).

## Data availability

Raw sequencing reads and statistical analysis have been deposited in the Gene Expression Omnibus (https://www.ncbi.nlm.nih.gov/geo/) under GEO accession number GSE262190.

## Acknowledgements

This study was supported by Fondation de la Recherche Médicale (FRM - DEQ20181039599), and the National Laboratory of Excellence program - Integrative Biology of Emerging Infectious Diseases (LabEx IBEID, ANR-10-LABX-62-IBEID).

## Competing interests

The authors declare no competing interests.

## FIGURE LEGENDS

**Supplementary Figure S1.**
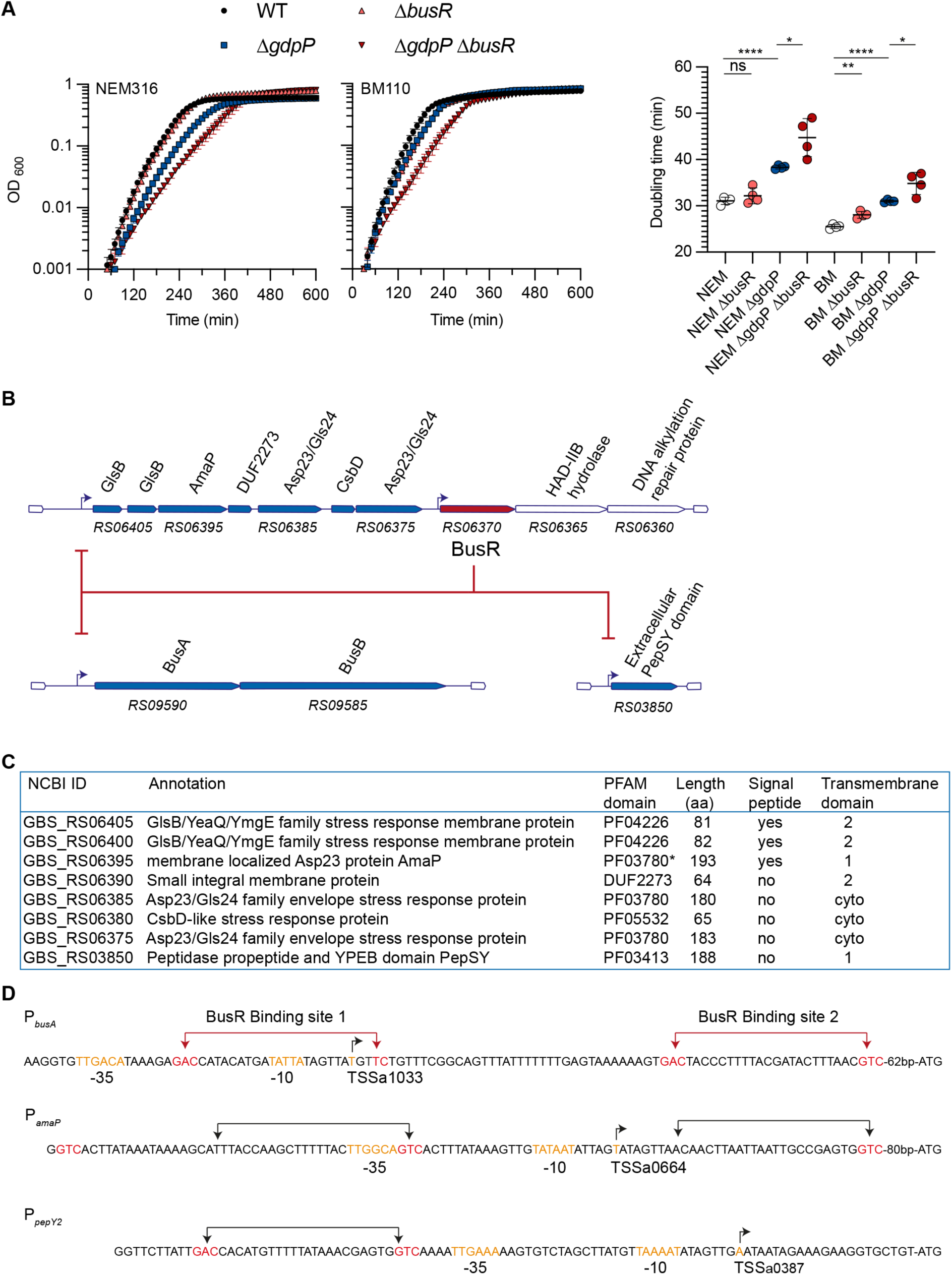
The BusR regulon. **(A)** Growth curves (left panel) and doubling time in exponential phase (right panel) of the WT strains (NEM316 and BM110), Δ*busR*, Δ*gdpP*, Δ*gdpP* Δ*busR* mutants in THY medium at 37°C. Means and SD are from a representative experiment with biological replicate (N = 4) analyzed using unpaired t-test (* p < 0.05, ** p < 0.01, **** p < 0.0001). **(B)** Schematic representation of the BusR regulon. Genes that are highly activated in the absence of BusR are represented by blue arrows, the relative length of the genes being conserved. NCBI gene ID are given in abbreviated form. (e.g., BusR = GBS_RS06370). **(C)** Genes annotation. The 7-genes *amaP*-operon encodes for AmaP, a membrane spanning protein with an Asp23-related domain (InterPro NCBIfam entry NF033218) predicted to act as a scaffold for the two cytoplasmic Asp23 proteins, and 4 uncharacterized small proteins. Signal peptides are predicted by Phobius and transmembrane domains by TMHMM integrated into the InterPro database. **(D)** Promoter sequences. The two experimentally characterized BusR binding sites on the P*_busA_* promoter are represented by red arrows with the canonical binding sites (GAC palindromes spaced 22 bp apart) and imperfect motif indicated with red letters. The GAC/GTC sequences in P*_amaP_* and P*_pepY2_* are also highlighted in red, with black arrows pointing 22 bp apart. The -35, -10, and transcriptional start site (TSS) are highlighted in orange.

**Supplementary Figure S2.**
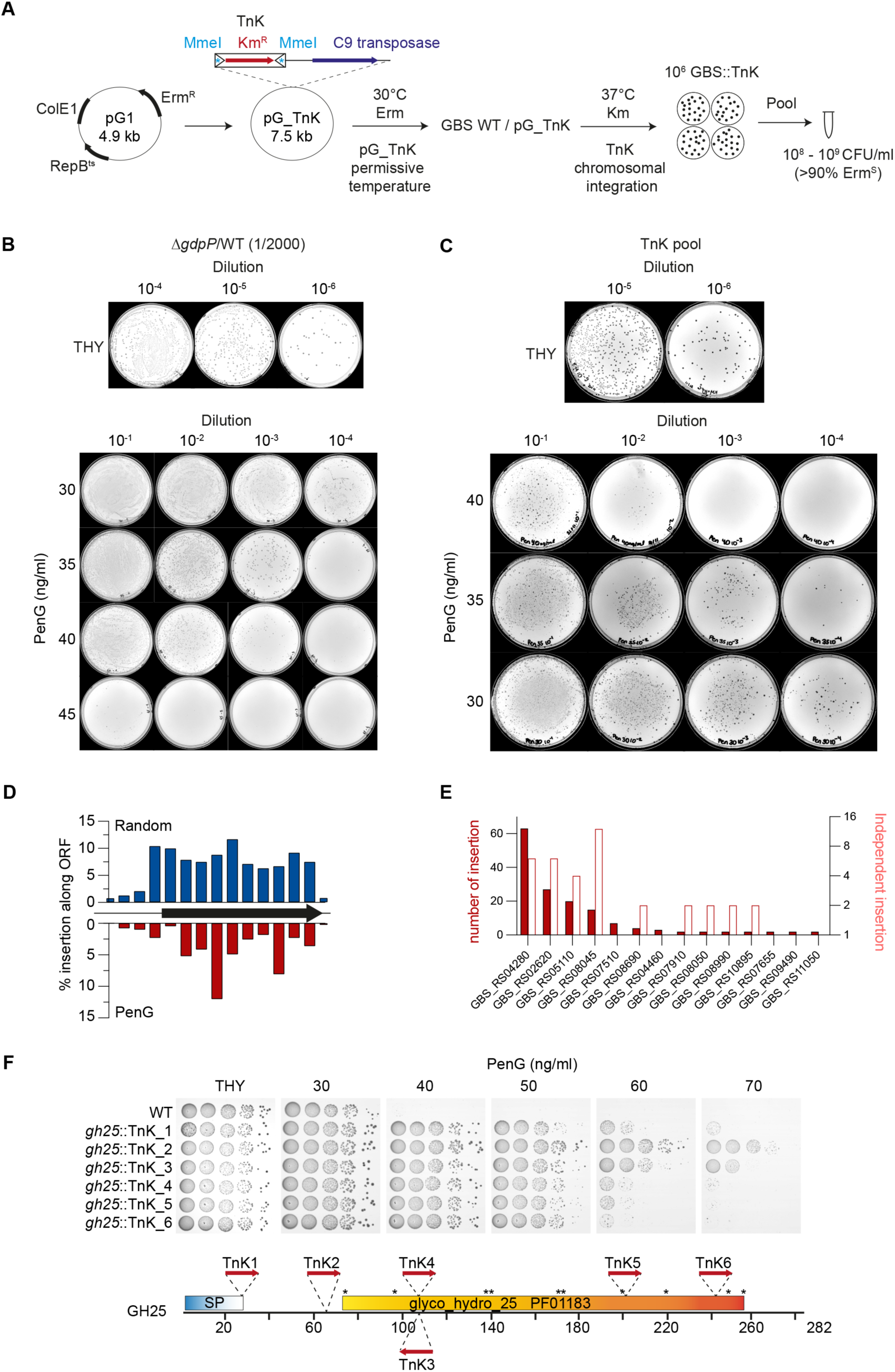
Genome-wide random screening for penicillin tolerance and reduced susceptibility. **(A)** Construction of the transposon library. The pG1 vector containing an erythromycin resistance gene (Erm^R^), a Gram-negative origin of replication (ColE1) and a thermosensitive Gram-positive origin of replication (RepB^ts^) was used as backbone. A cassette containing a custom mini-Himar transposon with a kanamycin resistance gene (red arrow) and MmeI restriction site in the inverted repeats was introduced in the pG1 altogether with a copy of the C9-transposase gene under a constitutive promoter. The pG_TnK plasmid was introduced in the NEM316 WT strain with a selection at 30°C (permissive temperature of replication) and erythromycin. After isolation, liquid cultures were diluted and plated on 160 kanamycin-containing plates. After incubation at 37°C (non-permissive temperature), approximately 10^6^ kanamycin resistant colonies were pooled, washed, and concentrated to 10^8^-10^9^ CFU/ml. Loss of the plasmid backbone were checked by numerating erythromycin versus kanamycin resistant colonies. **(B)** Screening condition for penicillin tolerance and reduced susceptibility. Cultures of the WT and Δ*gdpP* strains were mixed at a ratio of 2000 for 1 (V/V). The starting co-culture is diluted and spread on THY (upper plates) and penicillin-containing media (bottom plates). **(C)** Screening of the transposon library. Aliquots of the pool of 10^6^ insertional mutants were inoculated in THY, cultured for 1 hours at 37°C for recovery, and diluted before plating on THY (upper plates) and penicillin-containing media (bottom plates). Plates are incubated at 37°C for 24-36 hours in aerobic condition with 5% CO_2_. Colonies were then isolated on THY from the screening plates prior to genomic DNA extraction, transposon sequencing and phenotypic validation. **(D)** Distribution of transposon integration along ORFs. The length of the genes is normalized (horizontal black arrow) and subdivided into 10 bins. The proportion of transposon located in each bin is represent by vertical bars for mutants randomly isolated on THY (upper blue bars, N total= 240) and mutants isolated on penicillin-containing plates (lower red bars, N total = 191). For insertion outside ORFs, a binning of 50 base pairs was applied (4 bins from the start codon and 1 bin from the stop codon). **(E)** Genes frequently inactivated by transposon insertion recovered after penicillin screening. List of the 14 genes with the highest number of insertion (red bars, left axis) and independent chromosomal integration point (white bars, right axis) among a total of 191 mutants isolated on penicillin-containing plates. **(E)** Phenotype associated with independent transposon insertion into the GH25 glycose hydrolase. The highest number of insertions is in the gene *GBS_RS04280* encoding a 282 amino acids glycoside hydrolase belonging to the GH25 family (PF01183). Insertional mutants at 6 independent positions (red arrows: TnK1 to TnK6) were tested by spotting assay on penicillin-containing plates. SP: Signal peptide.

## SUPPLEMENTARY TABLES LEGENDS

**Supplementary Table S1: Transcriptomic analysis by RNA-seq.**

S1A: Δ*gdpP* and Δ*busR* mutants in the NEM316 WT strain - all genes. S1B: Δ*gdpP* and Δ*busR* mutants in the BM110 WT strain - all genes. S1C: Comparative analysis between NEM316 and BM110 homologues.

S1D: List of genes with significative p-adjusted value (< 0.0001) in at least one ΔgdpP mutant.

S1E: List of genes with significative p-adjusted value (< 0.0001) in at least one ΔbusR mutant.

**Supplementary Table S2: Strains used in this study and genome sequencing.**

S2A. Strain list with summary of sequenced genomes. S2B. NEM316 WT polymorphisms.

S2C. Unique polymorphisms in GBS mutants.

**Supplementary Table S3: Distribution of transposon integration into the NEM316 chromosome.**

S3A: Chromosomal coordinates of transposon integration in 240 colonies randomly isolated from THY.

S3B: Chromosomal coordinates of transposon integration in 191 colonies randomly isolated from penicillin plates.

S3C: Distribution of transposon integration by loci (PenG).

**Supplementary Table S4: Oligonucleotides and vectors used in this study.**

## Notes

### Competing Interest Statement

The authors have declared no competing interest.

